# Remdesivir Metabolite GS-441524 Effectively Inhibits SARS-CoV-2 Infection in Mice Models

**DOI:** 10.1101/2020.10.26.353300

**Authors:** Yingjun Li, Liu Cao, Ge Li, Feng Cong, Yunfeng Li, Jing Sun, Yinzhu Luo, Guijiang Chen, Guanguan Li, Ping Wang, Fan Xing, Yanxi Ji, Jincun Zhao, Yu Zhang, Deyin Guo, Xumu Zhang

**Author notes:** Correspondence to:Yu Zhang, Deyin Guo and Xumu Zhang. These authors contributed equally to this work (Yingjun Li, Liu Cao).

## Abstract

The outbreak of coronavirus disease 2019 (COVID-19) rapidly spreads across worldwide and becomes a global pandemic. Remdesivir is the only COVID-19 treatment approved by U.S. Food and Drug Administration (FDA); however, its effectiveness is still under questioning as raised by the results of a large WHO Solidarity Trial. Herein, we report that the parent nucleotide of remdesivir, GS-441524, potently inhibits the replication of severe acute respiratory syndrome coronavirus 2 (SARS-CoV-2) in Vero E6 and other cells. It exhibits good plasma distribution and longer half-life (t_1/2_=4.8h) in rat PK study. GS-441524 is highly efficacious against SARS-CoV-2 in AAV-hACE2 transduced mice and murine hepatitis virus (MHV) in mice, reducing the viral titers in CoV-attacked organs, without noticeable toxicity. Given that GS-441524 was the predominant metabolite of remdesivir in the plasma, the anti-COVID-19 effect of remdesivir may partly come from the effect of GS-441524. Our results also supported that GS-441524 as a promising and inexpensive drug candidate in the treatment of COVID-19 and future emerging CoVs diseases.

## Introduction

Coronavirus disease 2019 (COVID-19) caused by severe acute respiratory syndrome coronavirus 2 (SARS-CoV-2) was first identified in December 2019.^1, 2^ SARS-CoV-2 is a novel single-stranded RNA virus belonging to beta coronavirus.^3^ Although most coronavirus infections cause only mild respiratory symptoms, infection with SARS-CoV-2 could progress to acute severe respiratory syndrome (ARDS) and be lethal.^4^ Since the declaration of COVID-19 as a global pandemic by the World Health Organization (WHO), infection and mortality rapidly increased worldwide. As of 24 October 2020, more than 42,000,000 confirmed COVID-19 cases and >1,100,000 associated deaths were reported worldwide, according to the Johns Hopkins University COVID-19 global case dashboard. Effective therapies and drug candidates to treat COVID-19 are urgently needed.

Although several virus-based and host-based therapeutics agents have been evaluated for the treatment of COVID-19, such as lopinavir/ritonavir,^5^ immunoglobulin, hydroxychloroquine,^6^ EIDD-2801,^7^ baricitinib,^8^ and AT-527.^9^ Currently, only remdesivir, an RNA-dependent RNA polymerase (RdRP) inhibitor, recently received U.S. Food and Drug Administration (FDA) approval for COVID-19 treatment.^10, 11, 12, 13, 14^ However, its effectiveness to reduce hospital stay and risk of death of COVID-19 patients is still under questioning, as revealed by WHO solidarity trial.^15^ In addition, the broad applicability of remdesivir, especially accessibility in the developing country will be limited by its difficulty to synthesize and expensiveness. Structurally, remdesivir (Figure 1) is a nucleotide McGuigan prodrug presuming to be intracellularly metabolized into an active analog of triphosphate (GS-443902).^16^ The anionic phosphate moiety on remdesivir is masked by phenol and L-alaninate ethylbutyl ester, which is supposed to be enzymatically cleaved-off inside cells. However, pharmacokinetics (PK) study in nonhuman primates and healthy human subjects showed that remdesivir is rapidly metabolized to be GS-704277, GS-441524 (parent nucleoside), and GS-443902 in plasma following intravenous administration (Figure 1).^17, 18^ The major metabolite GS-441524 demonstrated a more persistent plasma exposure, a longer half-life (t_1/2_ ~ 27 h for GS-441524 vs. t_1/2_ ~ 1h for remdesivir) and a larger AUC value (AUC = 2230 hng/ml for GS-441524 vs. AUC = 1590 hng/ml for remdesivir).^19, 20^ The contribution of its metabolites especially GS-441524, to clinical outcomes is not fully understood.

**Figure1:**
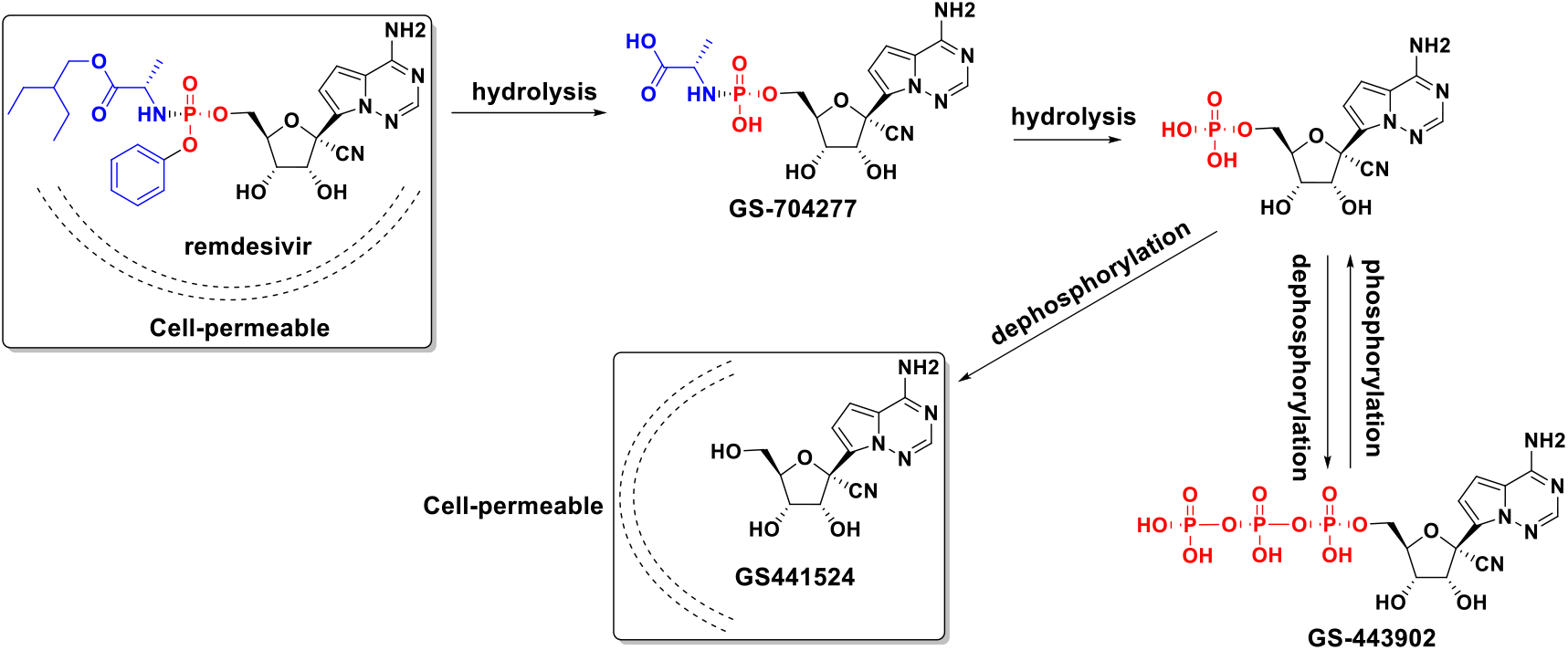
Proposed remdesivir metabolic pathway and chemical structures of metabolites.

*In vitro* studies revealed that GS-441524 was significantly less potent against Ebola virus (EBOV), hepatitis C virus (HCV) and respiratory syncytial virus (RSV), as compared to remdesivir.^12, 21^ The activation of nucleoside analogs needs two enzymatic phosphorylation steps to convert to the pharmacologically active triphosphate (GS-443902). The conversion of GS-441524 to monophosphate is considered a rate-limiting step in the activation of GS-441524. The prodrug remdesivir is considered to be superior to GS-441524 by bypassing the perceived rate-limiting first phosphorylation step and also by improving cell permeability.^22^ However, GS-441524 exhibited good *in vitro* efficacy against human coronaviruses (CoV), such as SARS-CoV, middle east respiratory syndrome coronavirus (MERS-CoV), and zoonotic coronaviruses. Feline infectious peritonitis (FIP) was caused by feline coronaviruses (FCoV) infection and has long been considered a fatal feline disease. A 96% cure rate was observed in GS-441524 treated cats with feline coronavirus infection.^16, 23, 24^ These early studies indicate the potential of GS-441524 for the treatment of coronavirus diseases as well as COVID-19.^25, 26^ Herein, we investigated and compared GS-441524 with remdesivir for their *in vitro* anti-SARS-CoV-2 activities, PK profile and *in vivo* efficacy against SARS-CoV-2 and MHV Our results provided further experimental insights to emphasize GS-441524 as a potential antiviral for COVID-19 and future emerging CoVs diseases.

## Results

### GS-441524 and remdesivir potently inhibit SARS-CoV-2 replication

Initially, we compared the anti-SARS-CoV-2 activity of GS-441524 and remdesivir in Vero E6 (African green monkey kidney cells), calu-3 (human lung adenocarcinoma cell) and caco-2 cells (colorectal adenocarcinoma). Cells were infected with SARS-CoV-2 at a multiplicity of infection (MOI) of 0.05 and treated with varying concentrations of remdesivir or GS-441524. Antiviral activities were evaluated by qRT-PCR quantification of viral copy number in the supernatant at 48 h post infection. GS-441524 and remdesivir potently inhibited SARS-CoV-2 replication in a dose-dependent manner, as showed in Figure 2. In Vero E6 cells, GS-441524 (IC_50_ = 0.70 μM) inhibited SRAR-COV-2 one-fold more potent than remdesivir (IC_50_ = 1.35 μM). The determination of intracellular viral RNA load in Vero E6 showed a similar trend to that in the supernatant (Figure 2D-E). In calu-3 and caco-2 cells, remdesivir exhibited better efficacy. The IC_50_ values of remdesivir in these two cells were 0.65 μM and 0.58 μM, and the IC_50_ values of GS-441524 in these two cells were 3.21 μM and 3.62 μM, respectively. Our results are consistent with the previously reported studies that the relative potencies of GS-441524 and remdesivir are cell type dependent.^25^ Cytotoxicity of the compounds in Vero E6, calu-3 and caco-2 cells were determined by the CCK8 assay. GS-441524 did not inhibit cell growth at high concentrations up to 50 μM, indicating its good safety profile (Figure 2A-C).

**Figure 2.**
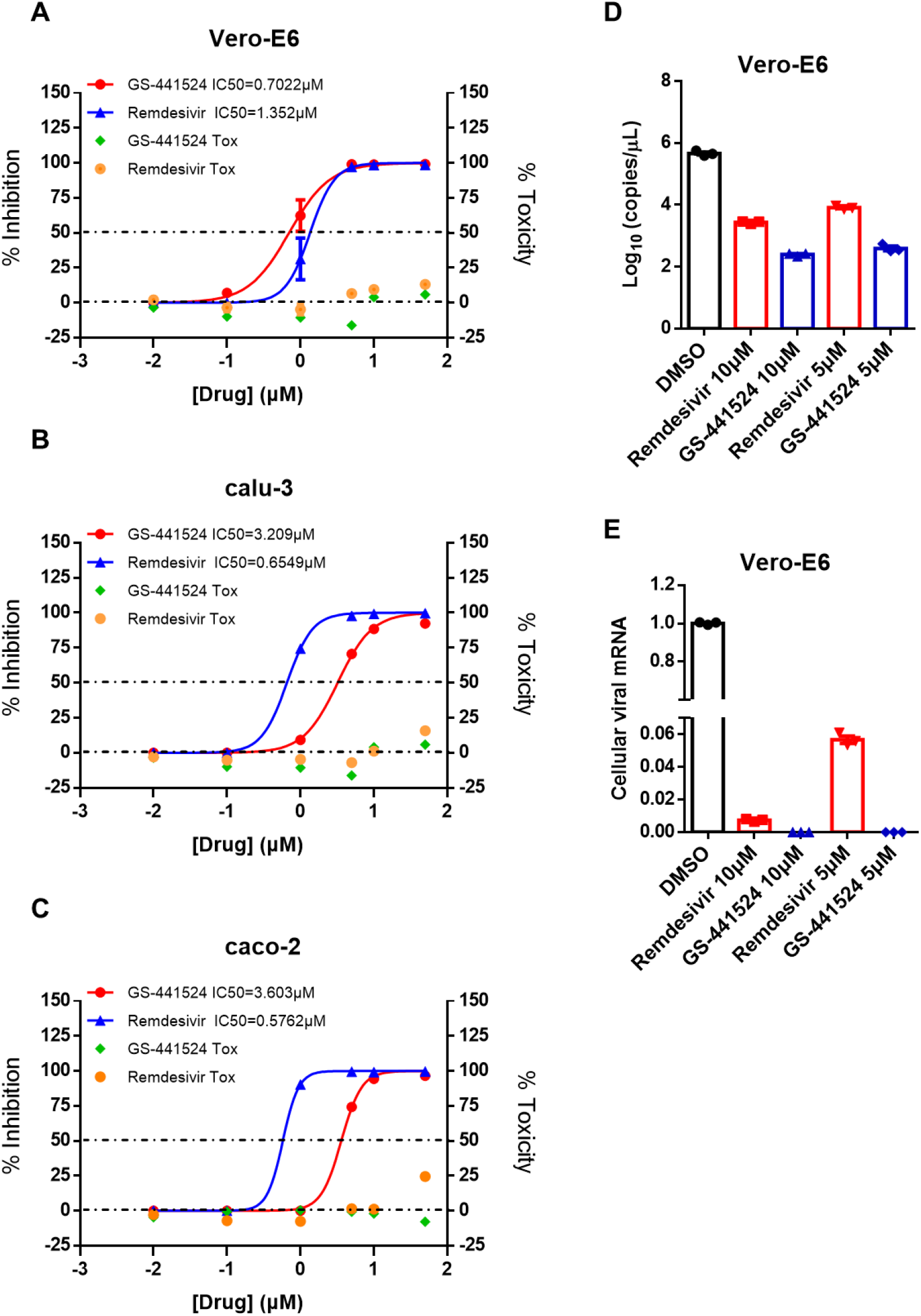
The prodrug remdesivir and parent nucleoside GS-441524 potently inhibit SARS-CoV-2 replication *in vitro*. Vero-E6 (**A**), calu-3 (**B**) and caco-2 (**C**) were infected with SARS-CoV-2 at an MOI of 0.05 and treated with different drugs (GS-441524 and Remdesivir) at different doses (0, 0.01, 0.1, 1, 5, 10, 50 μM) for 48 h. The viral yield in the cell supernatant was then quantified by qRT-PCR. Data represented are the mean value of % inhibition of SARS-CoV-2 on cells. At the same time, the cytotoxicity at different concentrations of drugs was tested. Vero-E6 cells were infected with SARS-CoV-2 at an MOI of 0.05 and treated with different doses (0, 5, 10 μM) of the indicated compounds for 48 h. The viral RNA in the cell supernatant (**D**) and intracellular (**E**) was then quantified by qRT-PCR.

### The PK profile of GS-441524 in rat

The PK profile of remdesivir and its metabolites was studied with intravenous (iv) administration of remdesivir in cynomolgus monkey and human subjects.^20, 27^ However, there is no reported PK about GS-441524 itself. Next, we determined the pharmacokinetics of GS-441524 in Sprague-Dawley (SD) rats *via* iv and intragastric (ig) injection with a dose of 30 mg/kg. GS-441524 exhibited encouraging iv PK parameters with a long half-life (t_1/2_) of 4.8 h and a high C_max_ of 163616.6 μg/L. Although the bioavailability (~5 %) of GS-441524 is not ideal, the C_max_ in ig administrated mice was 2708 μg/L (9.3μM), higher than the concentration required for > 50% SARS-CoV-2 inhibition, indicating that GS-441524 might be able to achieve in vivo efficacy relevant doses following an oral dosing regimen. Our results were consistent with the clinical PK study of remdesivir, which showed GS-441524, as a metabolite, has improved plasma stability as compared to that of remdesivir.

**Figure 3.**
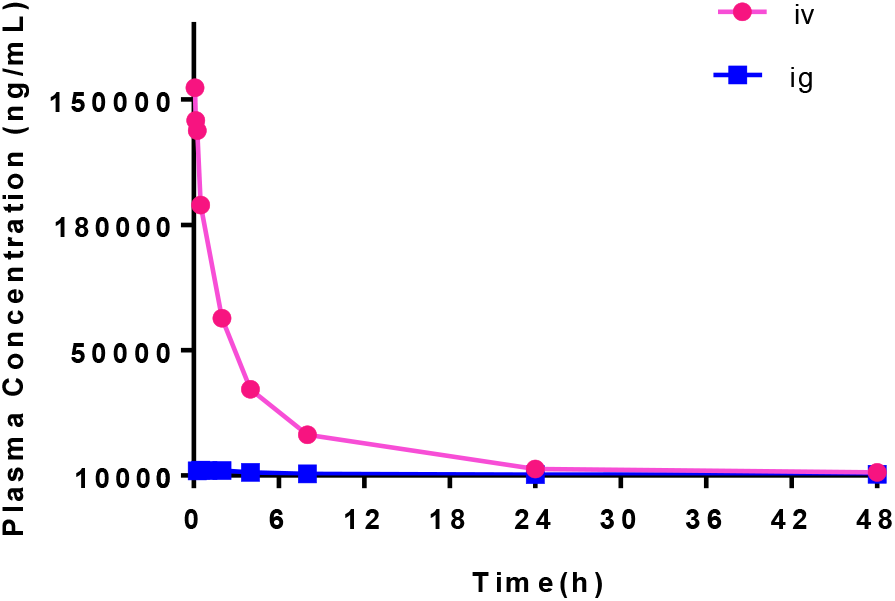
The time-concentration curve of GS-441524 in a PK study. **A.** plasma concentration and time curve following iv (red) and ig (blue) administration of 30 mg/kg GS-441524 in SD rat (data indicated are mean ± SD (standard deviation), n = 4).

**Table 1.** Pharmacokinetics parameters of GS-441524 in rat

|  | GS-441524 (iv) | GS-441524 (ig) |
| --- | --- | --- |
| AUC(0-t) ($\mu\text{g/L}\cdot\text{h}$) | 591916±265802 | 28668±7062 |
| t<sub>1/2</sub> (h) | 4.77±2.33 | 20.55±16.43 |
| T<sub>max</sub> (h) | 0.10±0.04 | 0.94±0.77 |
| C<sub>max</sub> ($\mu\text{g/L}$) | 163616.6±3747.3 | 2708.0±1308.8 |
| F % | | 4.84±1.19 |

### GS-441524 inhibits SARS-CoV-2 in AAV-hACE2 transduced mice

We next proceeded to study the *in vivo* anti-SARS-CoV-2 efficacy of GS-441524. Although mice are the convenient animal for *in vivo* assessing the anti-COVID-19 activity of drugs, they are resistant to SARS-CoV-2. SARS-CoV-2 uses human angiotensin-converting enzyme 2 (ACE2) to enter cells, but mouse ACE2 does not sensitize cells for infection.^28^ We established the transduction mice with adenovirus associated virus (AAV) vector expressing hACE2 in Biosafety Level 3 (BSL-3) laboratory. AAV-hACE2 mice support SARS-CoV-2 replication and exhibit pulmonary inflammation and lung injury.^29, 30^ In this study, 1×10^5^ plaque forming unit (PFU) SARS-CoV-2 were intranasally inoculated to AAV-hACE2 mice and mice body weights were monitored over a 10-day time course. Ip administration with GS-441524 one day prior to infection and continued dosing of 25 mg/kg/day for 8 days. SARS-CoV-2 infection of AAV-hACE2-sensitized mice were characterized by significant weight loss (more than 20%), severe pulmonary pathology and high-titer virus replication in the lung (Figure 4). GS-441524 demonstrated a significant virus clearance in lung at 2 dpi and protection of mice from body weight loss. Compared with mice treated with GS-441524, the lung tissues of untreated mice showed multiple injuries, including inflammatory cell infiltration from the trachea, peri-alveolar to interstitium. These data in sum were the first solid demonstration of the *in vivo* potency of GS-441524 against SARS-CoV-2.

**Figure 4.**
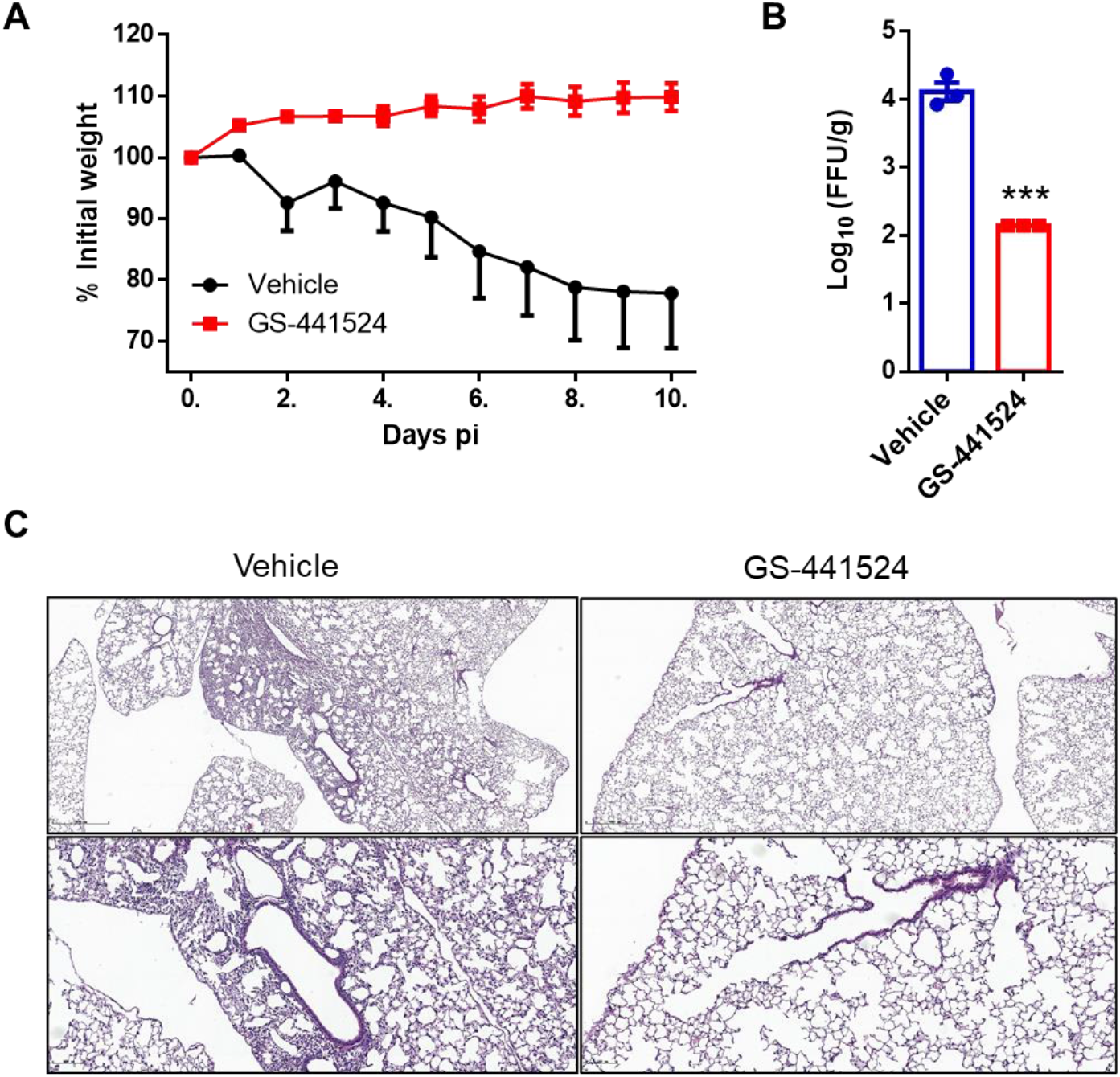
*In vivo* anti-SARS-CoV-2 efficacy of GS-441524 in mouse AAV-hACE2 model. AAV-hACE2 transduced mice were infected with SARA-CoV-2. The mice were administrated with either vehicle or GS-441524 25 mg/kg/day at −1 days pi (post innoculation) and the treatment was continued for a total of 8 days. Body weights were monitored every day (**A**). Lung tissues of 3 mice in each group were harvested and the viral titers were analyzed by qRT-PCR at 2 Dpi (**B**). (**C**) Representative Hematoxylin-eosin (HE) staining of lungs from hACE2 transduced mice Scale bars, 500mm (top) and 111 mm (bottom). *p values ≤ 0.05; **p values ≤ 0.005; ***p values ≤ 0.0005.

### The efficacy of GS-441524 in the treatment of mice MHV infection

MHV belongs to the coronavirus RNA viruses, sharing a common genus to SARS-CoV-2.^31^ MHV-A59 (A59 strain of MHV) infection caused hepatitis in mice, which could serve as a rapid experimental model to evaluate anti-CoVs agents. GS-441524 is known to be efficacious against MHV replication *in vitro* through RdRP inhibition.^29, 32^ However, whether it was able to reach the target organ and have *in vivo* effect was not known. To gain insight into the *in vivo* efficacy of GS-441524, mice were intranasally inoculated with MHV-A59 and the infection caused 87.5% of animal dead at 7 dpi (day post inoculation) (Figure 5A, group A). Administration of GS-441524 via ig (B1 group) or ip (intraperitoneal injection, B2 group) began at 0.5 hpi, with a dose of 100 mg/kg once (dose doubled) for the first day and 50 mg/kg/day for the following four days. The potent anti-MHV activities of GS-441524 was illustrated by the significant improvement of animal survival. (Figure 5A). The survival rate at the end of the experiment was 100% in group B1and B2 compared to 12.5% in group A.

**Figure 5.**
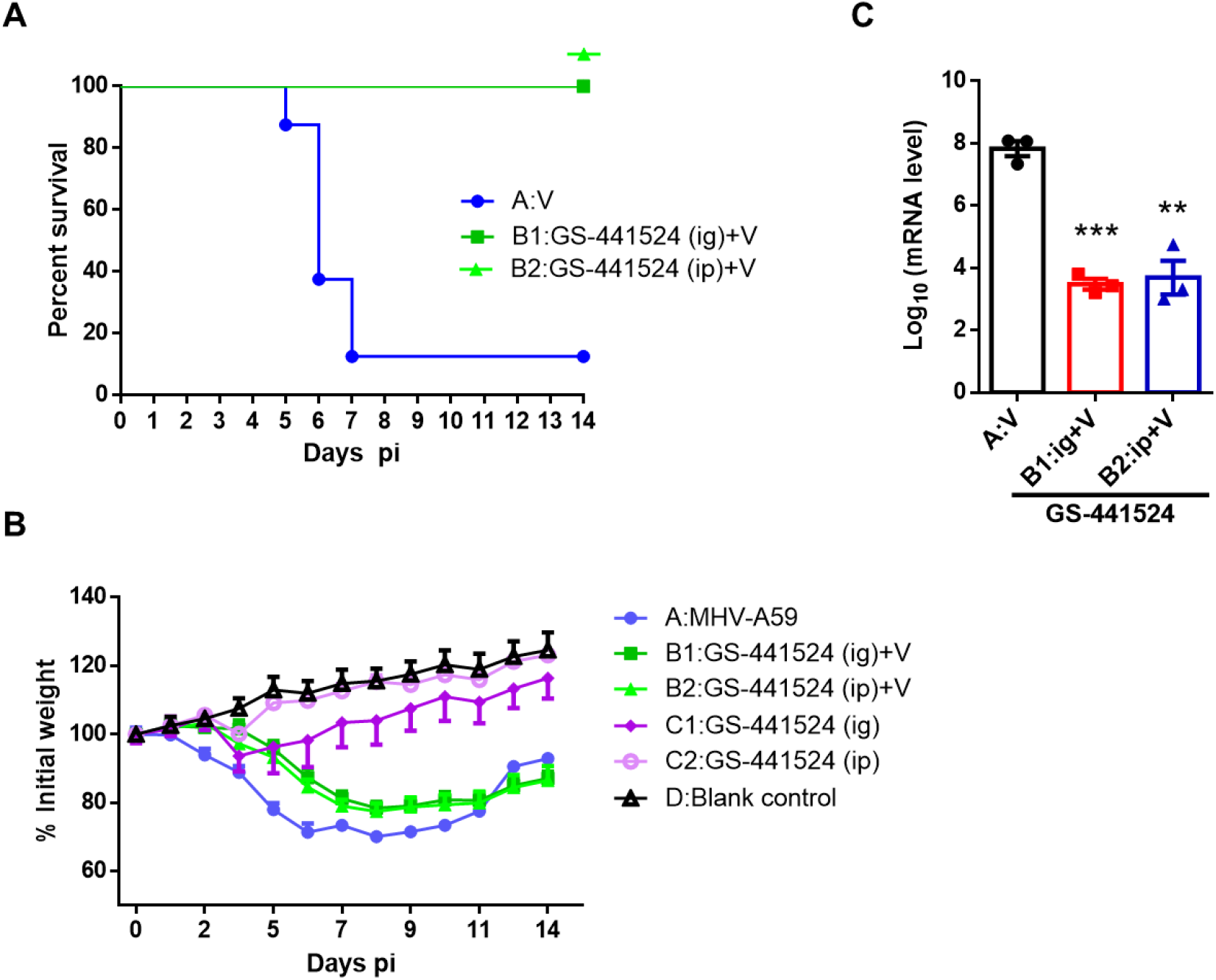
*In vivo* antiviral efficacy of GS-441524 in mice MHV-A59 model: mouse was randomly divided in 6 groups, MHV-A59 infected control (group A:V), GS-441524 50 mg/kg/day ig administration with MHV infection (group B1:GS-441524 (ig)+A), GS-441524 50mg/kg/day ip administration with MHV infection (group B2: GS-441524 (ip)+A), GS-441524 ig (group C1) and ip (group C2) control and blank control (group D). Drugs was treated for a continuous 5 days (dose doubled for the first day) beginning at 0.5h pi. Animal survival and body weight was showed in **A** and **B**. Liver tissues of 3 mice in either group A, B1or B2 were harvested and the viruses in the liver were analyzed by qRT-PCR (**C**) at 3 dpi. *p values ≤ 0.05; **p values ≤ 0.005; ***p values ≤ 0.0005.

To analysis the viral titration in the liver, RNA was extracted from lung tissue for qRT-PCR analysis at 3 dpi of the MHV infected mice (A, B1and B2 groups). GS-441524 treatment was able to significantly inhibit virus replication in the lung by more than 99.9% (Figure 5C). Moreover, drug treatment (C1 and C2 group) without virus inoculation did not significantly affect the body change, as compared to blank control (group D), indicating the excellent safety profile of GS-441524 (Figure 5B). The body weight loss of the virus inoculated mice (groups A, B1 and B2) might cause by the virus infection. These results were readily reproduced in a repeating experiment (**Figure S2**). Although the bioavailability of GS-441524 was low (about 5%), the current ig dosage was able to achieve *in vivo* therapeutic concentration and exhibited similar efficacy to other treatments, indicating the potential of GS-441524 as an oral drug.

## Discussion

The COVID-19 pandemic associated with high contagiousness, morbidity and mortality, emphasizing the imperative need for anti-viral agents. RdRP inhibitor remdesivir was the only anti-COVID-19 drug that accelerated the recovery in adults who were hospitalized with COVID-19 in a large randomized clinical trial, but controversy still remains.^15, 33^ Due to the synthesis complexity, expensiveness of Active Pharmaceutical Ingredient (API) and obligatory intravenous administration, the application and accessibility of remdesivir will be limited. The parent nucleotide GS-441524 is superior to remdesivir in several aspects. First of all, GS-441524 is a structurally simple molecule and is easier to synthesize than remdesivir. Secondly, pre-clinical and clinical PK study of remdesivir revealed that GS-441524 was the predominant and persistent metabolite in circulation. Although other metabolites, such as alanine metabolite GS-704277, triphosphorylated GS-443902 was also detected in the plasma, these compounds are highly negative charged and have poor cell membrane permeability, which will then inevitably convert to GS-441524 or excrete from the body (Figure 1). Thirdly, GS-441524 showed potent anti-viral activity against several stains of coronaviruses. However, it was reported that SARS-CoV-2 in cultured primary human airway epithelial (HAE) is much more sensitive to remdesivir than GS-441524 with a selectivity index of more than 1,000, questioning the efficacy of GS-441524 in lung tissue and COVID-19.^25^ Herein, we reported the *in vitro* and *in vivo* anti-SARS-CoV-2 activities of GS-441524. GS-441524 effectively inhibited the SARS-CoV-2 replication in all three cell lines (Vero E6, calu-3 and caco-2). It was able to rescue mice from death and reduce the viral load in the liver, either administrated ip, or ig. Importantly, GS-441524 showed significant efficacy against SARS-CoV-2 infection in mouse AAV-hACE2 model. We showed that GS-441524 potently inhibits SARS-CoV-2 replication in lung and alleviates lung inflammation and injury, suggesting the antiviral activity in human lung tissue and effectiveness against COVID-19 and highlighting the differences in *in vitro* HAE cultures and intact respiratory epithelial cells. Moreover, GS-441524 showed limited cytotoxicity and *in vivo* toxicity, demonstrating its remarkable safety profile together with the cytotoxicity screening as reported by Gilead and others.^12, 16^

The potent *in vivo* efficacy of GS-441524 against SARS-CoV-2 and MHV in mice model we reported here, together with the previous anti-FIP activity, demonstrate its broad application against zoonotic and human CoVs, its broad distribution and effectiveness to convert into the active GS-443902 across different tissues. Our results in sum indicated that part of remdesivir’s therapeutic effect resulted from the active metabolite GS441524 and the unnecessariness of the structural complexity of remdesivir. It was estimated that the pandemic of COVID-19 will be lasting for a long time. As zoonotic relative of CoVs repeatedly evolves to fatal CoVs, such as SARS-CoV, MERS-CoV and SARS-CoV-2, identification and evaluation of anti-viral therapies are urgently needed in the present and in the future. Our results supported GS-441524 is a promising drug candidate in the treatment of COVID-19 as well as future emerging coronavirus infection diseases, which might be helpful to address the current urgent need for safe and cheap antiviral treatment besides remdesivir.

## Methods

### Compounds, cells and viruses

Remdesivir (Cat. No. T7766) was purchased from TargetMol and GS-441524 (Cat. No. BCP35590) was purchased from Biochempartner. African green monkey kidney Vero E6 cell line (Vero-E6) was kindly provided by Dr. Hui Zhang (Sun Yat-sen University). Calu-3 and caco-2 were kindly provided by Dr. Yu Chen (Wuhan University). Vero-E6 and caco-2 were cultured in DMEM supplemented with 10% FBS, 100 U/mL penicillin and streptomycin at 37 °C in a humidified atmosphere of 5% CO_2_. Calu-3 cells were cultured in DMEM supplemented with 20% FBS, 100 U/mL penicillin and streptomycin, and 1% NEAA at 37 °C in a humidified atmosphere of 5% CO_2_. Patient-derived SARS-CoV-2 was isolated from a sputum sample from a woman admitted to the Eighth People’s Hospital of Guangzhou.^34^ MHV-A59 were obtained from the American Type Culture Collection. MHV-A59 were expanded in mouse liver cells NCTC 1469. Supernatants were collected and a passage 7 stock was subsequently stored at −80°C until used. SARS-CoV-2 infection experiments were performed in the BSL-3 laboratory of Sun Yat-sen University or Guangzhou Customs District Technology Center. MHV-A59 infection experiments were performed in the Biosafety Level 2 (BSL 2) laboratory of Guangdong Laboratory Animals Monitoring Institute.

### Antiviral activity assays

Vero E6, calu-3 and coco-2 cells were seeded at 1 × 10^5^ cells per well in 24-well plates. Cells were allowed to adhere for 16-24 h and then infected at MOI of 0.05 with SARS-CoV-2 for 1h at 37°C. Then viral inoculum was removed, and cells were washed 2 times with pre-warmed PBS. Medium containing dilutions of Remdesivir, GS-441524, or DMSO was added. At 48 hpi, supernatants or cells were harvested for qRT-PCR analysis. The dose-response curves were plotted from viral RNA copies versus the drug concentrations using GraphPad Prism 6 software.

### qRT-PCR analysis

For SARS-CoV-2 supernatant RNA quantification, RNA was isolated by E.Z.N.A.®Viral RNA Kit (OMEGA). SARS-CoV-2 nucleic acid detection kit (Daan Company) is used to detect the virus. For the detection of cellular viruses and tissue viruses, total RNA was isolated from cells or tissue samples with TRIzol reagent under the instruction of the manufacturer. The mRNAs were reverse transcribed into cDNA by PrimeScript RT reagent Kit (Takara). The cDNA was amplified by a fast two-step amplification program using ChamQ Universal SYBR qPCR Master Mix (Vazyme Biotech Co., Ltd). GAPDH was used to normalize the input samples via the ΔΔCt method. The relative mRNA expression level of each gene was normalized to GAPDH housekeeping gene expression in the untreated condition, and fold induction was calculated by the ΔΔCT method relative to those in untreated samples.

### CCK-8 cell viability assay

To investigate the effect of drugs on cell viability, Vero E6, calu-3 and coco-2 cells were seeded in 96-well plates at a density of 20,000 cells/well and were treated with drugs at indicated concentrations (0, 0.01, 0.1, 1, 5, 10, 50 μM) for 48 h. Cell viability was tested by using Cell Counting Kit-8 (CCK-8, Bimake, B34302). The figures were plotted from viral RNA copies in supernatants versus the drug concentrations using GraphPad Prism 6 software.

### PK analysis

SD rats (male, four animals per group) weighing 180–220 g were injected with GS-441514 intravenously (iv) and intragastricly (ig) at a dose of 30mg/kg. After administration, 0.3 mL of the orbital blood was taken at 0.083, 0.16 0.25, 0.5, 2, 3, 4, 8, 24 and 48 h for the iv group, and 0.25, 0.5, 1, 2, 3.0, 4, 6, 8, 24 and 48 h for the ig group, respectively. Samples were centrifuged under 4000 rpm/min for10 min at 4°C. The supernatants (plasma) were collected and stored at −20°C for future analysis. For plasma drug concentration analysis, an aliquot of 50 ul each plasma sample was treated with 100ul of 90% methanol and 600 ul of 50% acetonitrile mixture. The samples were centrifuged under 1200 rpm for 10min and filtered through 0.2 μm membrane filters. The drug concentration in each sample was tested by HPLC/MS. Analytes were separated on a InertSustain AQ-C18HP column (3.0 mm× 50 mm, 3.0μm, GL) using Waters UPLC/XEVO TQ-S. The pharmacokinetic parameters were calculated using DAS (Drug and Statistics) 3.0 software. The time-concentration curve was plotted using GraphPad Prism 6 software.

### AAV-HACE2 SARS-CoV-2 infected mice study

Adeno-associated virus 9 encoding hACE2 were purchased from Packgene (AAV-hACE2). The modified intratracheal aerosolization was used to intratracheally delivery AAV vector (5×10^11^ GC) to lung tissue of 4-week-old BALB/c mice.^35, 36^ Briefly, mice were anesthetized with 2.5% isoflurane in O_2_ (1 L/min). The 50 *μ*L AAV solution was then aerosolized at a rate of about 15 *μ*L/second using a microsprayer under fiberoptic laryngoscope. The mouse was further maintained anesthesia for 5 min to promote delivery of the vector deep into the lungs before transferring back to the cage. Thirty days post transduction, mice were intranasally infected with SARS-CoV-2 (1 × 10^5^ PFU) in a total volume of 50 *μ*L DMEM. Immediately, they were randomly divided into two groups (each group had 9 mice). A dosage of 25mg/kg/day of GS441524 or vehicle was ip administrated to the mice beginning at −1 dpi for a consecutive 8 days. Mice were monitored and weighed daily. 4 mice of each group were dissected at 2 dpi to collect lung tissues for virus titer detection and HE staining. All protocols were approved by the Animal Welfare Committee and all procedures used in this study complied with the guidelines and policies of the Animal Care and Use Committee.

### Focus forming assay (FFA)

The viral titration in lung tissue was determined using focus FFA assay as previously described.^30^ Vero E6 cells were seeded in 96-well plates one day before infection. Lung homogenates were serially diluted and used to inoculate Vero E6 cells at 37°C for 1 h. Inocula were then removed before adding 125 μL 1.6% carboxymethylcellulose per well and warmed to 37°C. After 24 h, cells were fixed with 4% paraformaldehyde and permeabilized with 0.2% Triton X-100. Cells were then incubated with a rabbit anti-SARS-CoV-2 nucleocapsid protein polyclonal antibody (Cat. No.: 40143-T62, Sino Biological), followed by an HRP-labeled goat anti-rabbit secondary antibody (Cat. No.: 109-035-088, Jackson ImmunoResearch Laboratories). The foci were visualized by TrueBlue Peroxidase Substrate (KPL), and counted with an ELISPOT reader (Cellular Technology). Viral titers were calculated as per gram tissue.

### Hematoxylin and Eosin (HE) Staining

Mice lung dissections were fixed in zinc formalin and embedded with paraffin. Tissue sections (~4 μm) were stained with hematoxylin and eosin.

### Mouse MHV efficacy study

3 to 4 weeks specific-pathogen-free (SPF) male BALB/c (Guangdong Medical Experimental Animal Center) were maintained in microisolated cages and housed in the animal colony at the biosafety level 2 facility at Guangdong Laboratory Animals Monitoring Institute. Mice were fed standard lab chow diet and water ad libitum. The mice were randomly divided into eight groups and anesthetized by respiratory with isoflurane. Immediately, Mice in Group A, B1 and B2 (each group have 15 mice) received an intranasal inoculation of 30μL MHV-A59 (TCID_50_ =10^-7.125^/100μL) in PBS. Control groups (Group C1, C2 and D, each group have 3 mice) were mock treated with phosphate-buffered saline (PBS). Mice in B1 and B2 group received GS-441524 via ig (Group B1) or ip (Group B2) at 0.5h pi at a dose of 100 mg/kg respectively and mice in A group received PBS instead. Mice in control groups (Group C1, C2 and D) received GS-441524 or PBS in the same way. GS-441524 were administered 50 mg/kg once a day in the following four days. Group A and D administered 0.2 mL of PBS. Mice were monitored daily for symptoms of disease: including body weights, clinical symptoms and death for14 days. 4 mice of group A, B1 and B2 were dissected at 3 dpi to collect liver tissues for virus titration detection. All protocols were approved by the Animal Welfare Committee.

### Statistical analysis

All values are mean ± SD or SEM of individual samples. Data analysis was performed with GraphPad Prism Software (GraphPad Software Inc., version 6.01). The statistical tests utilized are two-tailed and respective details have been indicated in figure legends. *P* values of < 0.05 were considered statistically significant. (*, *P* values of ≤ 0.05. **, *P* values of ≤ 0.005. ***, *P* values of ≤ 0.0005. ****, *P* values of ≤ 0.0001).

## Acknowledgment

The project was supported by Shenzhen Science and Technology Innovation Committee (ZDSYS20190902093215877), Shenzhen Bay Laboratory (SZBL2019062801006) and Technology and National Natural Science Foundation of China (grant #32041002). We thank Chuwen Lin from School of Medicine, Sun Yat-Sen University for the help in lung pathology analysis.

## Author contributions

X.Z. and D.G. initiated the project; X.Z., D.G., Y.Z. and Y.L. designed the project. Y.L. and L.C. wrote the manuscript. Y.L., G.L., and P.W. prepared the compounds; L.C., F.X. and Y.J. performed the antiviral activity experiments; G.L., F.C., J.S., Y-Z.L. and J.Z. carried out the MHV mice experiments; Y-F.L. performed the PK study. YL and L.C. analyzed the data. Y.Z., D.G and X.Z. supervised and supported the project. All authors reviewed and approved the manuscript.

## Additional information

Supplementary information and chemical compound information are available in the online version of the paper. Reprints and permissions information is available online at www.nature.com/reprints. Correspondence and requests for materials should be addressed to Y.Z., D.G. or to X. Z.

